# An architectural role of *oskar* mRNA in granule assembly

**DOI:** 10.1101/2023.08.31.555701

**Authors:** Mainak Bose, Branislava Rankovic, Julia Mahamid, Anne Ephrussi

## Abstract

Ribonucleoprotein (RNP) granules are membraneless condensates that organize the intracellular space by compartmentalization of specific RNAs and proteins^1^. Studies have shown that RNA tunes the phase behavior of RNA binding proteins (RBPs)^2–4^, but the role of intermolecular RNA-RNA interactions in assembly of RNP granules *in vivo* remains less explored^5–7^. Here, we determine the role of a sequence-specific RNA-RNA kissing-loop interaction in assembly of mesoscale *oskar* RNP granules in the female *Drosophila* germline. A two-nucleotide mutation that disrupts kissing-loop-mediated *oskar* mRNA dimerization impairs condensate formation *in vitro*, *oskar* granule assembly in the developing oocyte - leading to defective posterior localization of the RNA, and abrogation of *oskar*-associated processing bodies (P-bodies) upon nutritional stress. This specific *trans* RNA-RNA interaction acts synergistically with the scaffold RBP, Bruno^8^, in driving condensate assembly. Our study highlights the architectural contribution of an mRNA and its specific secondary structure and tertiary interactions in formation of an RNP granule essential for embryonic development.

---

Eukaryotic cells contain a large number of RNA-protein condensates, broadly known as RNP granules, including stress granules, germ granules, neuronal transport granules and others^1^. These membraneless compartments are enriched in RNAs and RBPs, many of which harbor intrinsically disordered regions (IDRs) and prion-like domains (PrLDs)^4^. Recent studies on biomolecular condensation have highlighted the role of RNA-RBP interactions in regulating the assembly of RNP granules^1,2^; here, the synergistic action of stereospecific RNA-RBP and multivalent IDR-IDR interactions drives the formation of multicomponent condensates, which have also been described using the theoretical “scaffold-client” framework^9,10^. A generic model of RNP granule formation postulates that RNA binding can promote high local concentration of IDR-containing RBPs, which through protein-protein interactions connect individual RNP complexes into mesoscale condensates. What is commonly neglected in such models is the potential role of intermolecular RNA-RNA interactions. Long RNA molecules can form higher-order assemblies also in the absence of proteins^11^. This is exemplified by protein-free condensation of RNA homopolymers *in vitro*^12^, cell-free total yeast RNA^7^ and pathogenic repeat-expansion RNAs^5^. The nature and biophysical properties of intermolecular RNA-RNA interactions span a continuum from high-affinity sequence-specific interactions to promiscuous base-pairing between exposed sequences along large RNAs^1,13^.

*oskar* granules in the *Drosophila* female germline are a class of transport RNPs that package and localize the maternal RNA *oskar* to the posterior of the developing oocyte. The locally translated Oskar protein is essential for abdominal patterning and specification of germ cell fate during embryonic development^14,15^. We have recently reported that *oskar* transport granules are phase-separated condensates with solid-like material properties^8^. Using a combination of *in vitro* and *in vivo* assays, we identified the RBP Bruno as a primary scaffold protein that is crucial for *oskar* granule formation and their liquid-to-solid phase transition. The RNA Recognition Motifs (RRMs) of Bruno bind specific sequences (Bruno Response Elements/BREs) in the *oskar* 3’ untranslated region (UTR)^16^. Bruno N-terminal PrLD self-association drives assembly of mesoscale *oskar* granules, which partition client proteins in an RNA-dependent manner to regulate diverse aspects of *oskar* function, such as translation regulation^8^. Fluorescence microscopy-based quantifications estimated an average of 16 *oskar* mRNA (3 kb) molecules packaged in ∼400 nm diameter condensates with an RNA concentration of 873 nM^8^. The high RNA concentration suggests a potential architectural role of *oskar* mRNA in granule assembly or organization. In fact, our minimal *in vitro* reconstitutions with *oskar* and Bruno indicated formation and stabilization of RNA-RNA interactions upon Bruno-driven condensation^8^. However, the nature of *trans* RNA-RNA interactions and their contribution to *oskar* granule condensation remained to be explored.

## An RNA kissing-loop interaction is critical for *oskar* granule assembly

The 3’UTR of *oskar* mRNA harbors a 67 nucleotide long stem-loop structure, SL2b, also referred to as the Oocyte Entry Signal (OES) (Fig. 1a). The AU-rich stem of SL2b serves as a *cis*-acting RNA localization signal essential for Dynein-dependent transport of *oskar* from the nurse cells to the oocyte^17^. The terminal loop of the same stem-loop structure harbors a six nucleotide palindromic sequence (5’ CCGCGG 3’) known to promote dimerization of *oskar* mRNA *in vitro* via a kissing-loop interaction through canonical Watson-Crick base pairing^18^ (Fig. 1b). Dimerization of the OES is robust *in vitro* and occurs in absence of any monovalent or divalent cations in the buffer. Aiming to disrupt the kissing-loop intermolecular interactions, we introduced a two nucleotide substitution (5’ <u>UU</u>GCGG 3’) in the palindrome, hereon referred to as *oskar* UU^18^. This indeed abolished OES dimer formation even under conditions with high concentrations of Na^+^ and Mg^2+^ ions (Fig. 1b). We previously reported that, *in vivo*, the kissing-loop-based dimerization promotes co-packaging of transgenic reporter RNAs with endogenous *oskar* RNA^18,19^. This observation suggests a role of the kissing-loop in assembly of endogenous *oskar* granules. To test this hypothesis, we expressed genomic *oskar* transgenes comprising either a wild type (*oskar* WT) or a mutated dimerization domain (*oskar* UU) in flies in which no endogenous *oskar* RNA is expressed. In absence of *oskar*, oogenesis fails to progress^20,21^. Both the transgenic WT and UU RNAs enriched in the early oocyte and rescued progression of oogenesis in the *oskar* RNA null flies (Fig. 1c). Therefore, the two-nucleotide substitution in the loop does not disrupt the recruitment of the Dynein-transport machinery to the OES stem^17^, suggesting that the UU mutation does not interfere with the secondary structure of the stem loop. From mid-oogenesis onwards, however, the UU RNA mislocalized along the oocyte cortex and frequently accumulated as a cluster near the posterior pole, whereas the WT RNA robustly localized at the posterior pole (Fig. 1c, d). The *oskar* UU transport phenotype was not due to defects in oocyte polarity, as evident from the antero-lateral position of the oocyte nucleus at stage 9 and the proper localization of *gurken* mRNA (Extended Data Fig. 1). Careful examination revealed a diffuse distribution of the *oskar* RNA signal in the *oskar* UU oocytes, in contrast to the distinct, granular signal in the *oskar* WT, suggesting a defect in *oskar* granule formation in the mutant (Fig. 1e). Quantification of the RNA signal showed a significant reduction in partitioning of *oskar* UU RNA into granules, confirming that loss of the kissing-loop interaction interferes with granule assembly *in vivo* (Fig. 1f). We previously identified Bruno, a translation repressor of *oskar*, as a scaffold protein that drives *oskar* granule assembly *in vivo* in the presence of wild type *oskar* mRNA^8^. The present observations suggest that in addition to Bruno, the *oskar* mRNA itself might play a structural role in scaffolding the granules *in vivo*.

**Fig. 1:**
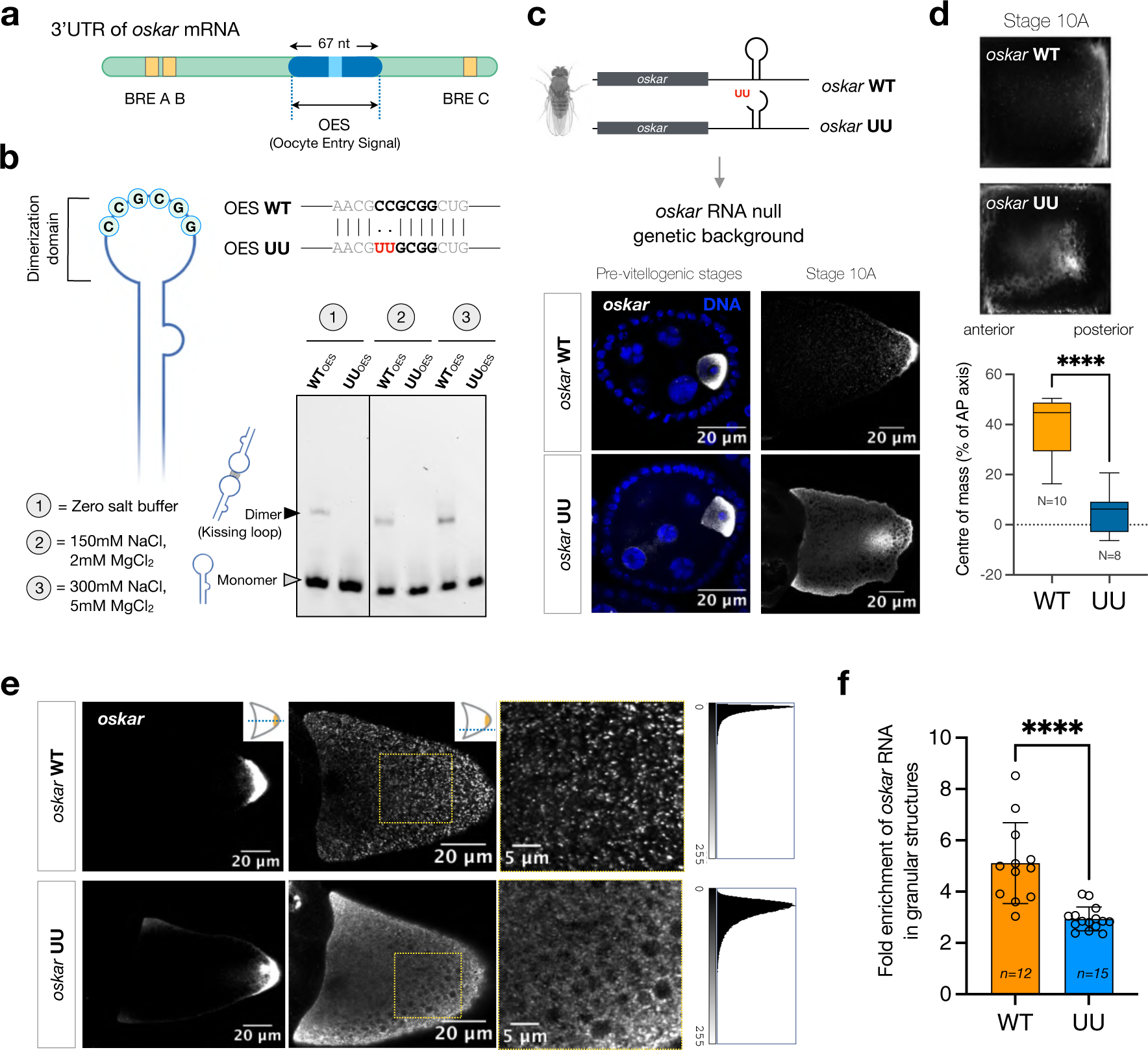
A kissing-loop RNA-RNA interaction is essential for *oskar* granule assembly *in vivo*. **a**, Schematic representation of the *oskar* 3’UTR showing the relative positions of the OES and the BREs. **b**, Cartoon representation of the 67 nucleotide-long OES highlighting the palindromic sequence that engages in a kissing-loop interaction with another *oskar* molecule. Dimerization of the WT and UU mutant OES in buffers with varying salt concentrations using atto633-labeled *in vitro* transcribed RNAs. **c**, Localization of *oskar* mRNA (grayscale) in pre-vitellogenic and stage 10 egg chambers detected by single molecule FISH (smFISH) in *oskar* WT and *oskar* UU transgenic flies. Experiments were carried out in flies genetically null for endogenous *oskar* RNA. **d**, Average *oskar* RNA signal (grayscale) from multiple stage 10 oocytes, anterior to posterior. Plot shows quantification of the data; position of the *oskar* center of mass relative to the geometric center of the oocyte (dotted horizontal line) along the anteroposterior (AP) axis. **e**, Representative confocal images of *oskar* RNA (grayscale) in the *oskar* WT and *oskar* UU oocytes at the equatorial (right) and cortical (middle) planes of the egg chamber. Right panels are enlarged views of the cortical plane images (yellow box). Histograms of pixel intensities of the enlarged area reveal the diffuse, non-punctate signal of *oskar* RNA in the case of *oskar* UU. **f**, Intensity-based segmentation of the granules from the cytoplasm was performed and enrichment of *oskar* RNA in the granules quantified. Error bars represent SD, and n denotes the number of oocytes analyzed. Unpaired Student’s t-tests were used for comparisons. Significance level: **** < 0.0001.

## The kissing-loop acts as a specific RNA ‘sticker’

To understand the mechanistic contribution of the kissing-loop interaction in scaffolding granules, we truncated the *oskar* 3’UTR to include the last 359 nucleotides, which harbor the OES and the 3’-most 166 nucleotides of the RNA that are crucial for oogenesis progression^21^. The *in vitro* transcribed WT_359_ RNA was predominantly dimeric and also oligomerized into multiple higher-order species, as evident from their slower mobility on a native gel in the electrophoretic mobility shift assay (EMSA; Fig. 2a). In contrast, the UU_359_ RNA exhibited significantly less dimerization. Since the mutant OES alone failed to dimerize *in vitro* (Fig. 1b), the observed dimeric form of UU_359_ potentially arises from promiscuous RNA-RNA contacts. Indeed, a dimeric species was still detectable when the entire stem-loop was deleted from the 359 nucleotide long RNA (*oskar*_292_, Fig. 2b), confirming that additional RNA-RNA interactions were promoted by sequences outside the stem-loop structure. However, when subjected to a thermal gradient, the dimeric form of the WT_359_ was more resistant to denaturation than that of the UU_359_ or *oskar*_292_ (Fig. 2b), confirming that in contrast to promiscuous, weak RNA-RNA interactions, the sequence-specific kissing-loop interaction stabilizes the RNA dimer. Notably, the higher-order species observed in WT_359_ persisted in stringent buffer conditions, indicating that the WT_359_ RNA can robustly self-assemble by virtue of the kissing-loop as well as additional RNA-RNA interactions promoted by the remainder of the RNA sequence (Fig. 2b, c). Upon heating, these higher-order oligomers melted faster than the kissing-loop-induced dimer, further highlighting the strength of the kissing-loop interaction in stabilizing RNA tertiary contacts (Fig. 2b). Therefore, the kissing-loop acts as a specific RNA ‘sticker’ in initiating RNA multimerization, which is propagated by additional non-specific intermolecular RNA ‘stickers’, while the flanking ‘spacer’ RNA sequences contribute to expanding the network, potentially by providing binding sites for RBPs^22,23^ (Fig. 2c).

**Fig. 2:**
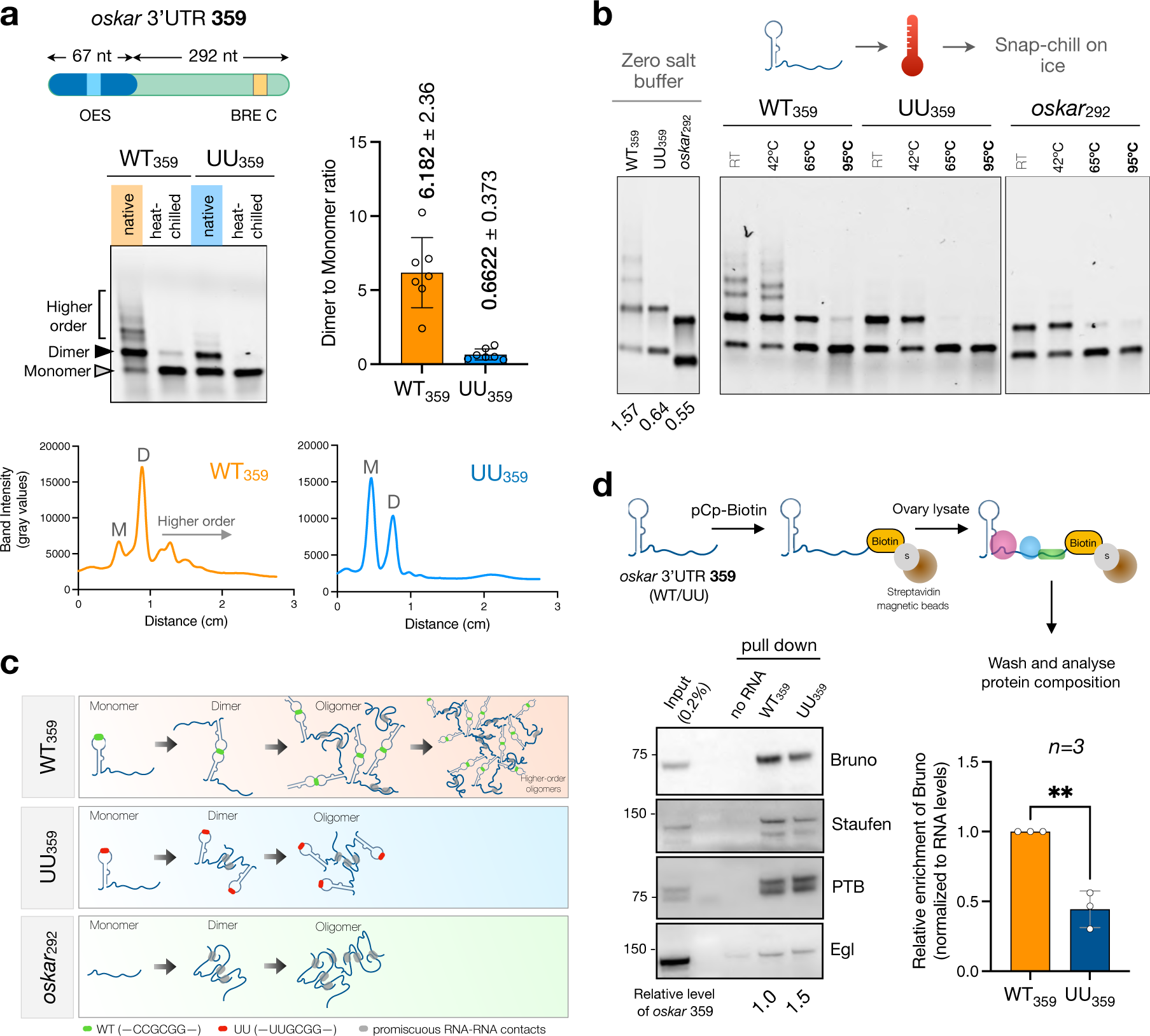
Mutation in the kissing-loop reduces association with scaffold protein Bruno. **a**, Schematic representation of the 359 nucleotide fragment of *oskar* 3’UTR; the 67 and 292 nucleotide sub-fragments are indicated. Electrophoresis of the atto633-labeled WT_359_ and UU_359_ RNAs under native and heat-chilled conditions in EMSA buffer (containing 150 mM NaCl, 2 mM MgCl_2_) shows enhanced formation of dimeric and higher-order oligomeric species by the WT_359_ RNA. Lane intensity profiles of the RNA signal under native conditions are plotted below. The graph on the right represents the dimer-to-monomer ratio quantified from multiple electrophoresis experiments. **b**, Behavior of the WT_359_, UU_359_ and *oskar*_292_ RNA fragments under stringent buffer conditions in absence of Na^+^ and Mg^2+^ ions (left) shows that both the OES mutant (UU_359_) as well the OES-deleted (*oskar*_292_) RNAs form dimers *in vitro*. The dimer to monomer ratio for each condition is denoted below the gel. WT_359_, UU_359_ and *oskar*_292_ RNAs were incubated in EMSA buffer at the indicated temperatures followed by snap-chilling on ice and subsequent native agarose gel electrophoresis. **c**, Graphical model illustrates the transition from monomeric to oligomeric species of the indicated RNAs. Note that owing to the degenerative nature of RNA-RNA base pairing, promiscuous intermolecular interactions (gray ovals) are prevalent for all three RNAs. **d**, Schematic workflow of the RNA affinity capture experiment performed using 3’-biotinylated WT_359_ and UU_359_ RNAs and ovary lysate from wild type *Oregon-R* flies. Beads without RNA serve as a negative control. Representative western blot data showing differential association of bona fide *oskar* granule RBPs. The relative level of the respective RNAs (determined by qPCR) pulled down by the streptavidin beads are indicated below the blot. Quantification of the Bruno levels was carried out using western blots from three independent replicates after normalization with the amount of pulled-down RNA. Error bars represent SD, and n denotes the number of oocytes analyzed. Unpaired Student’s t-tests were used for comparisons. Significance level: ** < 0.01.

Next, we tested if the UU mutation affects association of bona fide RBPs with the RNA. Transcript-specific isolation of RNPs demands a large amount of starting material and is experimentally challenging in the case of *Drosophila* oocytes^24^. Therefore we resorted to an *ex vivo* approach, whereby we biotinylated the synthesized WT_359_/UU_359_ RNA at the 3’ end, immobilized the RNA on magnetic streptavidin beads and incubated it with ovary extracts from wild type flies (Fig. 2d). Bona fide *oskar* RNA binding proteins such as Bruno^16^, Staufen^25^ and PTB^26^ were specifically captured using this affinity pull down method. Interestingly, the amount of the core scaffold protein Bruno associated with UU_359_ was significantly lower than with the WT_359_ RNA (Fig. 2d and Extended Data Fig. 2). Since both the WT_359_ and UU_359_ RNAs contain the BRE C, differential recruitment of Bruno suggests an impact of RNA dimerization on Bruno association.

## The kissing-loop interaction is crucial for condensation with Bruno

To investigate if Bruno is differentially associated with the *oskar* WT and UU RNAs in the egg chamber, we used EGFP knock-in (KI) Bruno flies^27^ in which the WT and UU transgenes were expressed in absence of endogenous *oskar* RNA. In the cytoplasm of the nurse cells, Bruno-EGFP signal colocalized with *oskar* RNA on microtubule tracks^8^ for both *oskar* WT and *oskar* UU (Fig. 3a), indicating that the UU mutation does not abrogate Bruno binding *in vivo*. However, in the oocyte, as opposed to the granular signal and strong colocalization observed with *oskar* WT, the Bruno-EGFP signal was largely diffuse and rarely colocalized with *oskar* RNA signal in the case of *oskar* UU (Fig. 3a). Considering our *in vitro* results which show the importance of kissing-loop-driven higher-order oligomerization for Bruno association, we hypothesized that mutation of the kissing-loop does not impair binding of the scaffold protein Bruno, but hinders higher-order oligomerization of the Bruno-RNA complex, thus aborting granule assembly *in vivo*. To test this, we performed EMSAs with Bruno-EGFP and *oskar*. Although both the WT_359_ and UU_359_ RNAs bound Bruno, the WT_359_ formed higher-order RNP complexes to a significantly greater extent than the UU_359_ RNA (Fig. 3b). A similar difference in higher-order complex formation was observed with the full length *oskar* 3’UTR (Extended Data Fig. 3a, b). EMSA with the OES alone did not show any detectable Bruno-binding (Fig. 3c), confirming that the stem-loop does not associate with Bruno and suggesting that Bruno binding is restricted to the remaining 292 nucleotide sequence (*oskar*_292_) which harbors the BRE C. Indeed, the *oskar*_292_ RNA bound Bruno and formed higher-order oligomers, showing that Bruno binding/oligomerization can be uncoupled from RNA dimerization (Fig. 3c). Since the 292 nucleotide sequence is identical in the WT_359_ and UU_359_ RNAs, this observation strongly suggests that in addition to Bruno-driven oligomerization, the kissing-loop interaction contributes to higher-order species formation.

**Fig. 3.**
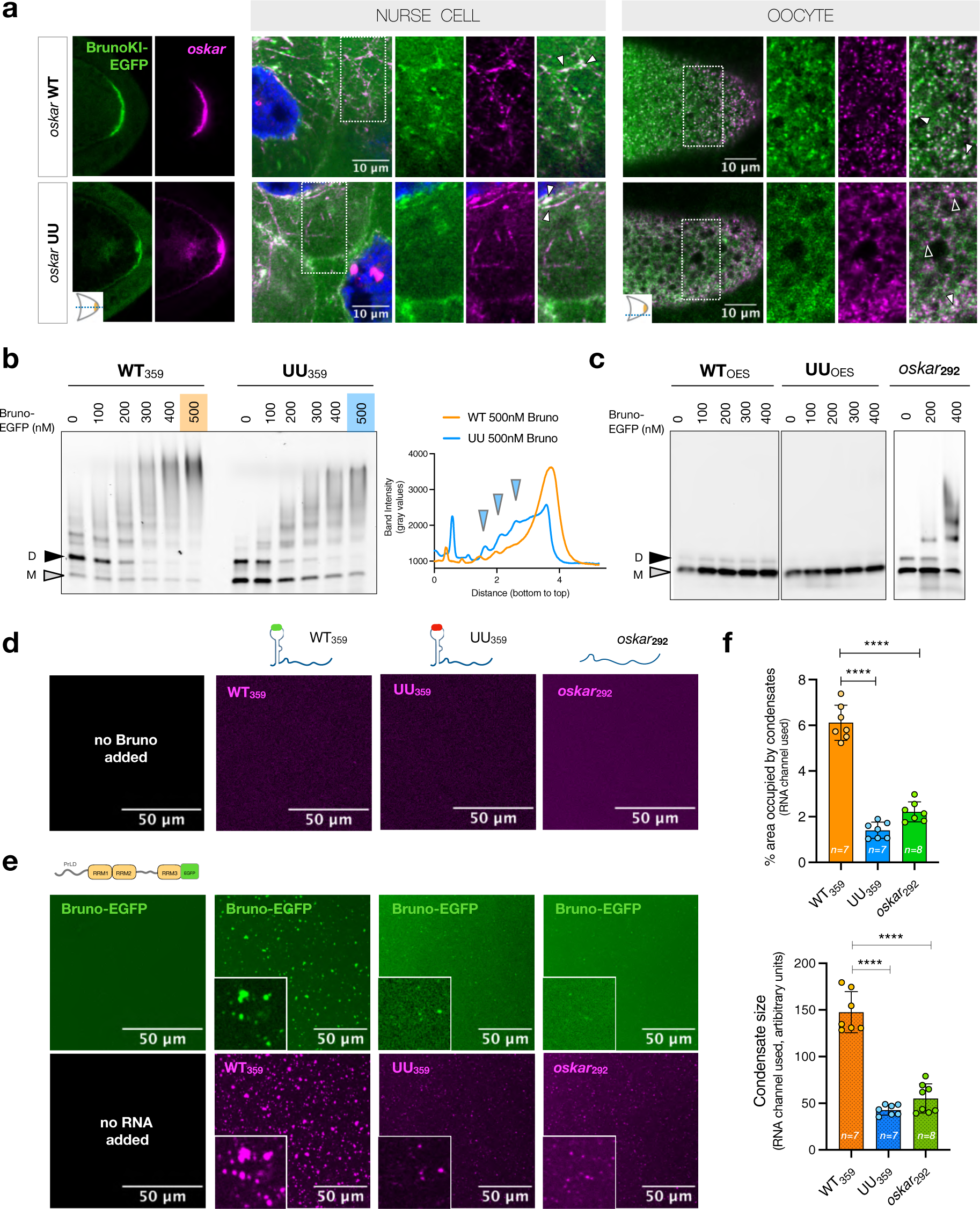
The kissing-loop is essential for condensate assembly with Bruno. **a**, Localization of Bruno-EGFP (green) and *oskar* WT or UU RNA (magenta) *in vivo*. Snapshots of the oocyte posterior (equatorial plane) show enrichment of Bruno KI-EGFP at sites where *oskar* RNA is highly concentrated. 1 µm thick confocal slice of nurse cell and oocyte (cortical plane); boxed areas enlarged on the right. Filled white arrowheads: colocalization of Bruno and RNA; empty arrowheads: lack of association of *oskar* RNA signal with Bruno. **b**, EMSA using 50 nM atto633-labeled WT_359_ and UU_359_ RNAs with indicated concentrations of Bruno-EGFP. Lane intensity profiles (from bottom to top of the gel) at the 500 nM Bruno concentration are plotted on the right. Blue arrowheads indicate intermediate oligomeric complexes detectable only in the case of UU_359._ **c**, EMSA using 50 nM atto633-labeled WT_OES_, UU_OES_ and *oskar*_292_ RNAs with indicated concentrations of Bruno-EGFP showing that RNA dimerization can be uncoupled from Bruno-driven oligomerization. **d-e**, *in vitro* phase separation assay using 5 µM Bruno-EGFP (green) and 100 nM of the respective atto633-labeled RNAs (magenta). **f**, Quantification of condensate size and area occupied (based on the RNA channel) are plotted. Error bars represent SD, and n denotes the number of oocytes analyzed. Unpaired Student’s t-tests were used for comparisons. Significance level: **** < 0.0001.

Given the established role of Bruno as a granule scaffold^8^, we next investigated the significance of RNA-RNA interactions in the context of Bruno-driven condensate assembly *in vitro*. The experiments were carried out using Bruno below the saturation concentration (C_sat_), where Bruno condensation is not detected. The RNA fragments also did not form any visible assemblies on their own under the chosen close-to-physiological buffer conditions with 150 mM NaCl and devoid of crowding agents (Fig. 3d). However, mixing the protein and the RNA led to spontaneous co-condensation of Bruno and WT_359_ into microscopic condensates, whereas the UU_359_ and Bruno formed few assemblies that were significantly smaller in size (Fig. 3e, f). Interestingly, the *oskar*_292_ RNA behaved identically to the UU_359_ despite binding and oligomerization with Bruno (Fig. 3e, f). Multivalent interactions between protein and RNA chains are the driving force for RNP condensate network formation. Our experiments show that the intermolecular kissing-loop RNA-RNA interaction, as well as Bruno-driven oligomerization, co-scaffold *oskar* condensate assembly (Fig. 4). Therefore, in the case of *oskar* granules, multivalency encoded by the PrLD-PrLD interactions acts synergistically with the intermolecular kissing-loop interaction to establish the *oskar* ribonucleoprotein network. Interfering with either Bruno phase separation as we have previously shown by deletion of the PrLD^8^, or disrupting the RNA kissing-loop interaction (this study) is detrimental to *oskar* granule formation. These RNP assemblies provide the platform for recruitment of additional effector proteins that regulate *oskar*’s functions in germline development.

**Fig. 4.**
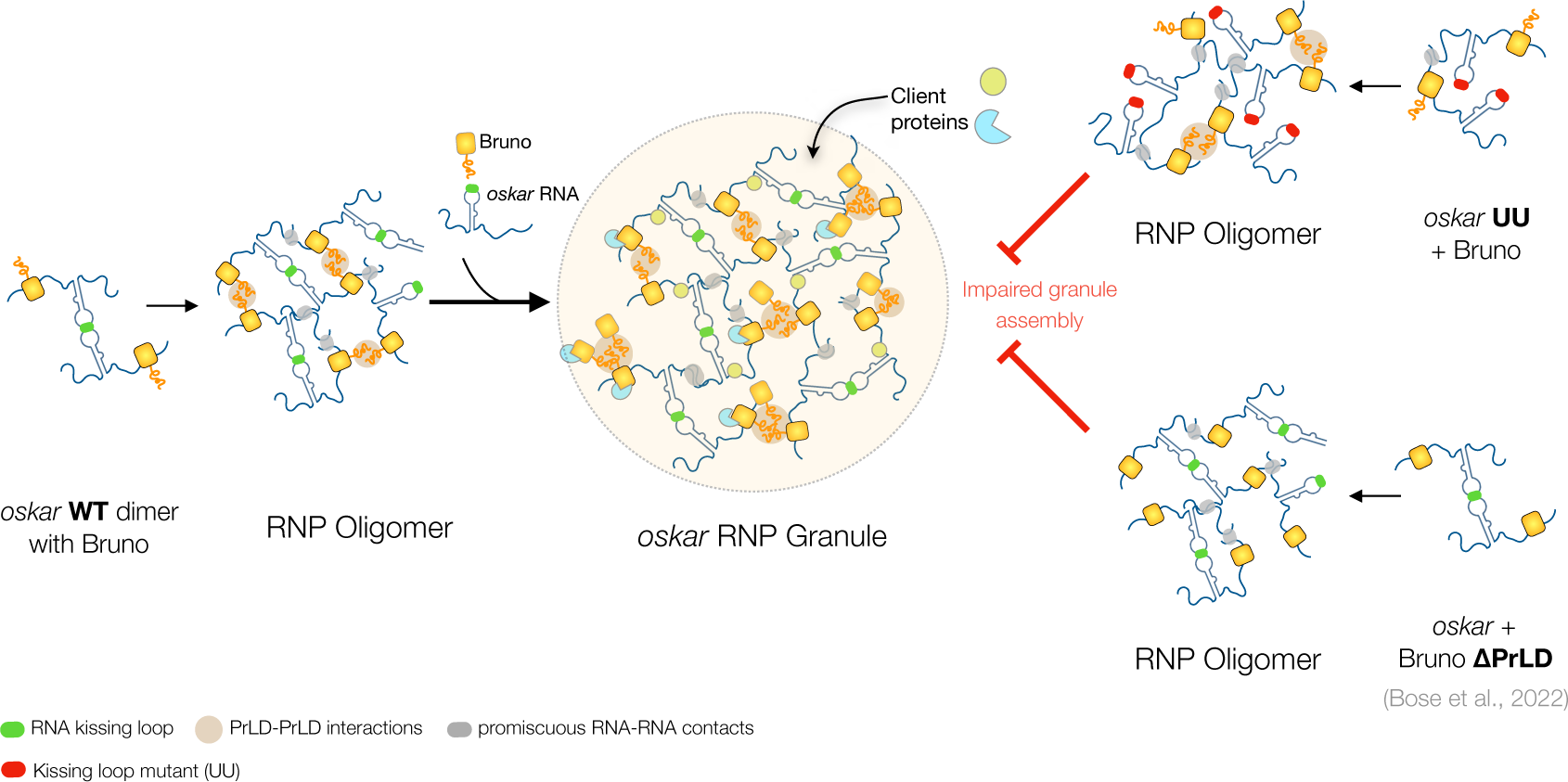
A combinatorial model of *oskar* RNP granule assembly. Schematic model showing the assembly of *oskar* granules as a result of cooperative multivalent interactions between the scaffold molecules *oskar* and Bruno. The model depicts how the combinatorial action of RNA-RNA (RNA kissing-loop and promiscuous inter-RNA contacts), RNA-protein (sequence-specific binding of Bruno to the *oskar* 3’UTR) and protein-protein (Bruno PrLD-PrLD) interactions drive oligomerization and phase separation of *oskar* RNP granules.

## The kissing-loop interaction is essential for P-body formation in the germline

Another class of RNP assemblies containing *oskar* mRNA is observed in the *Drosophila* female germline specifically upon nutrient deprivation^28,29^ (Fig. 5a, b and Extended Data Fig 4a). These stress-induced assemblies are referred to as P-bodies^28^, as they share protein components such as Me31B/Dhh1/DDX6 with yeast and mammalian P-bodies^30^. The localization of *oskar* mRNA to P-bodies is specific, as endogenous *bicoid* and *gurken* mRNAs were not enriched in these stress-induced assemblies (Extended data Fig. 4b). We observed that in addition to endogenous *oskar*, Bruno also localized to P-bodies (Fig. 5b). Furthermore, while the transgenic *oskar* WT RNA localized to P-bodies in absence of endogenous *oskar* RNA (Extended data Fig 4c), more than 75% of egg chambers expressing *oskar* UU failed to form P-bodies after 6 h of nutrient deprivation (Fig. 5c, d). Our observations indicate that *oskar* mRNA is an integral and essential component of these assemblies. While a mechanistic understanding of how these large RNPs form under conditions of nutritional stress is lacking, the fact that a two-nucleotide substitution in the kissing-loop of *oskar* disrupts P-body formation strongly suggests that, in addition to its architectural role in *oskar* transport granules, this high affinity sequence-specific mRNA interaction mode also has a key role in P-body formation, and highlights the importance of RNA-RNA interaction in driving diverse higher-order RNP assemblies *in vivo*.

**Fig. 5.**
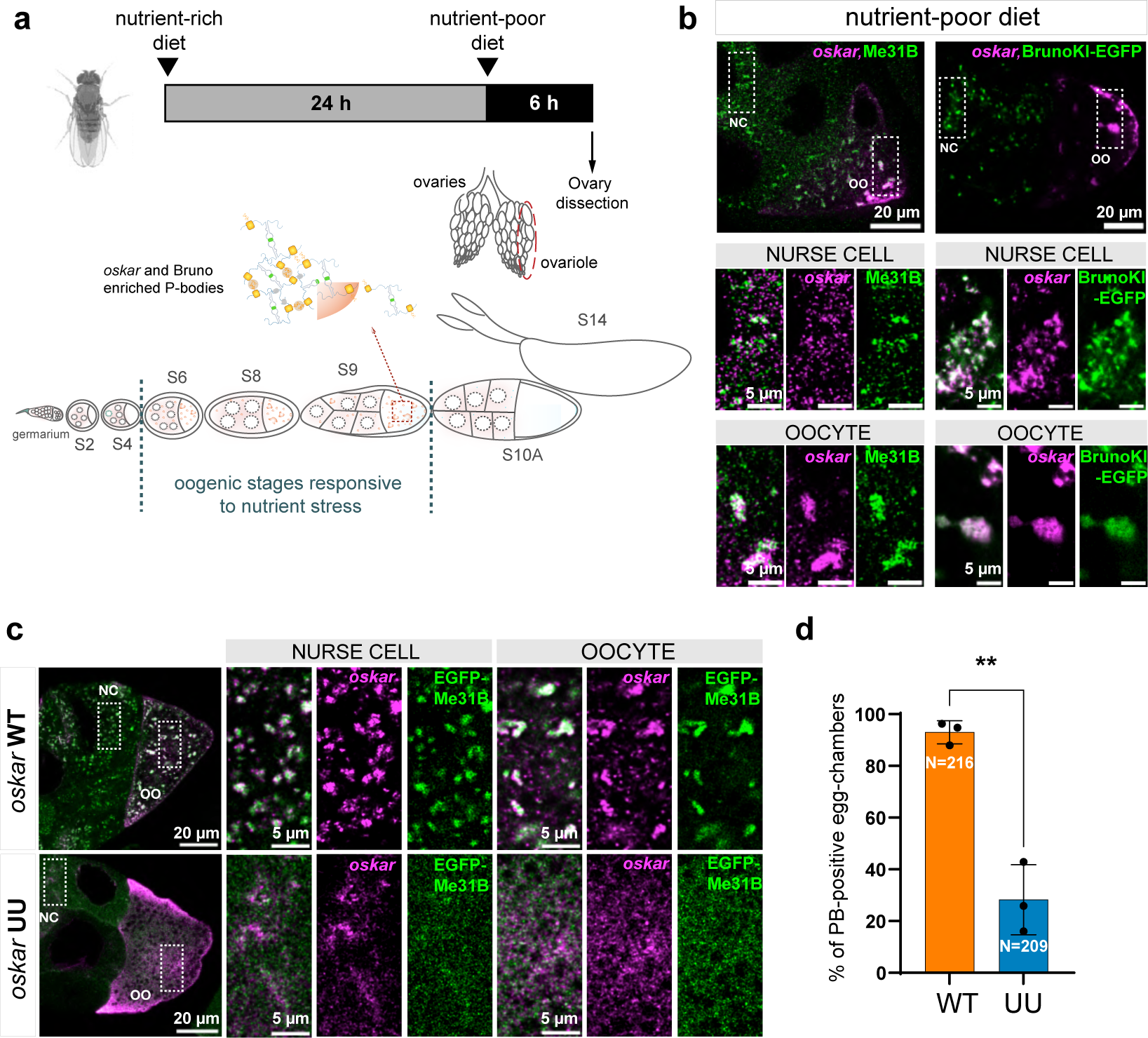
Mutation in the kissing-loop impairs nutritional stress-induced P-body formation. **a,** Schematic of the experimental procedure for nutrient deprivation of female flies and of a *Drosophila* ovariole showing the oogenesis stages analyzed for P-body formation in response to nutrient stress. **b,** Co-detection of *oskar* mRNA (smFISH; magenta) and Me31B (immunofluorescence; green) in wild-type (*w*^1118^) egg chambers (left panels) and *oskar* RNA smFISH (magenta) and Bruno-EGFP (green) in BrunoKI-EGFP egg chambers (right panels), after 6 hours of nutrient deprivation. **c-d,** Representative confocal images of *oskar* RNA (magenta) and EGFP-Me31B (green) in mid-oogenesis stage egg chambers of *oskar* WT and *oskar* UU flies after 6 h of nutrient deprivation. Boxed areas are enlarged on the right. Percentage of egg chambers forming P-bodies (PB-positive) in *oskar WT* and *oskar UU* flies. Three biological replicates were analyzed and egg chambers at stages 6-9 were scored. Experiments were carried out in flies genetically null for endogenous oskar RNA. For the statistical analysis unpaired two-tailed t-test was used (**, p ≤ 0.01) and mean ±SD is plotted. Note that to allow better visualization of the fluorescent signal in the nurse cell compartment, brightness and contrast were adjusted during image processing.

## Conclusion

Our findings demonstrate that a long mRNA molecule does not merely act as a passive scaffold that concentrates RBPs by sequence-specific binding to promote multivalent protein-protein interactions that drive granule assembly. Rather, sequence-specific *trans* RNA-RNA interactions contribute to the granule network formation. The primary scaffold RBP, Bruno, binds to both wild-type and mutant RNA, but is insufficient to drive condensate assembly in absence of the kissing-loop interaction. RNA kissing-loop interactions are well studied in retroviruses, whose genome is a dimer of two genomic RNAs (gRNA) of positive polarity^31^. Dimerization of the gRNA is highly conserved and essential for the viral life-cycle^32^. In the case of HIV-1, the Dimerization Initiation Site (DIS) comprises a six nucleotide GC-rich palindrome which nucleates the intermolecular tertiary contact between two copies of the gRNA via a kissing-loop interaction (loose dimer) followed by a switch to an extended duplex (tight dimer) that stabilizes the dimer^33,34^. However, for *oskar* mRNA, a switch to an extended conformation does not occur in the oocyte, as evident from the predicted OES structure *in vivo* inferred from mutational profiling and sequencing of dimethyl sulfate modified RNAs (DMS-MapSeq)^35^. Additionally, super-resolution STORM imaging using FISH probes spanning the entire length of *oskar* mRNA revealed a consistent drop in signal radius when probing near the stem loop compared to the rest of the RNA sequence, confirming sequence-specific *trans* RNA-RNA interactions being predominant in this region of the 3’UTR^36^.

Specific intermolecular RNA-RNA interactions can stem from canonical Watson-Crick base pairing between complementary RNA stretches and from Hoogsteen-type base pairing of G-rich tracts that form four-stranded G-quadruplexes, as reported for RNAs associated with repeat expansion disorders^5^. Specific RNA-RNA contacts also arise from kissing-loop interactions, which involve only very short palindromic sequences that can nevertheless impart exceptional mechanical stability to RNA dimers, as observed with artificial hairpins^37^. In fact, due to their cohesive nature, kissing-loops are often engineered into RNAs to promote intermolecular contacts and facilitate three dimensional packing of RNAs for high resolution structural studies^38,39^. In addition to sequence-specific interactions, non-specific contacts are also highly prevalent in long RNA fragments, as seen in our *in vitro* experiments using transcribed RNAs. Inside the cell, such promiscuous interactions presumably occur at high local concentrations of RNA, such as upon stress-induced release of mRNAs from polysomes^22^, in transcription foci^40^ or in RNA-rich condensates such as paraspeckles, which harbor up to 50 copies of the 23 kb long *NEAT1_2* long non-coding RNA inside a ∼300 nm condensate^41,42^. However, in our experiments, the promiscuous interactions not only failed to stabilize the dimeric species when exposed to thermal fluctuations, but also could not drive condensate formation *in vitro* and *in vivo* upon the addition of the RBP. It is likely that within *oskar* granules, the prevalence of such non-sequence specific RNA-RNA interactions depends on the availability of protein-free RNA segments. In contrast, the kissing-loop-mediated intermolecular interaction is crucial and acts as a specific RNA ‘sticker’^23^, and together with Bruno PrLD-PrLD interactions, helps in physical crosslinking of the RNAs into an interconnected network with solid-like physical properties^8^, which is reinforced by additional promiscuous RNA-RNA contacts. Such a nucleated RNA-protein meshwork can facilitate partitioning of client proteins that can regulate condensate functions. The advantage of sequence-specific RNA-RNA interactions in condensation is to dictate compositional specificity with respect to the RNA species. This is critical for *oskar* granule function, as co-mixing with other maternal transcripts is detrimental for embryonic development^43,44^. Evidence of such RNA species-specific condensates exists in the filamentous fungus *Ashbya gossypii* where RNA sequence and secondary structure specifies condensate RNA composition^6^. In fact, in the *Drosophila* oocyte, a similar kissing-loop interaction leads to dimerization of *bicoid* mRNA, the maternal anterior determinant, which facilitates RNP formation with the dsRNA binding protein Staufen^45^.

In summary, we demonstrate that a specific intermolecular RNA-RNA interaction in cooperation with an intrinsically disordered RBP can scaffold the assembly of a RNA-protein condensate that functions in germline specification and embryonic body axis patterning in *Drosophila*.

## Acknowledgements

We thank Imre Gaspar for sharing his observation that *oskar* UU and wild-type flies respond differently to nutrient deprivation and providing R-scripts for visualization of particle segmentation data. We are grateful to Akira Nakamura for providing the EGFP-Bruno flies. We acknowledge Pravin Jagtap, Janosch Hennig and Frank Wippich for insightful discussions. M.B. was supported by DFG-FOR 2333 grant EP 37/4-1 from the Deutsche Forschungsgemeinschaft (Germany) to A.E. and by the EMBL, B.R. was supported by a studentship of the Darwin Trust of Edinburgh and the EMBL. J.M. and A.E. acknowledge funding from the EMBL.

## Author Contributions

All authors contributed to the design of the study, interpreted the results and wrote the manuscript. M.B. performed the *in vitro* experiments and analyzed the data. B.R. established the conditions for nutrient deprivation of flies and harvested ovary samples for biochemical analysis. M.B. and B.R. carried out imaging and analysis of *Drosophila* egg chambers.

## Extended Data

**Extended Data Fig. 1.**
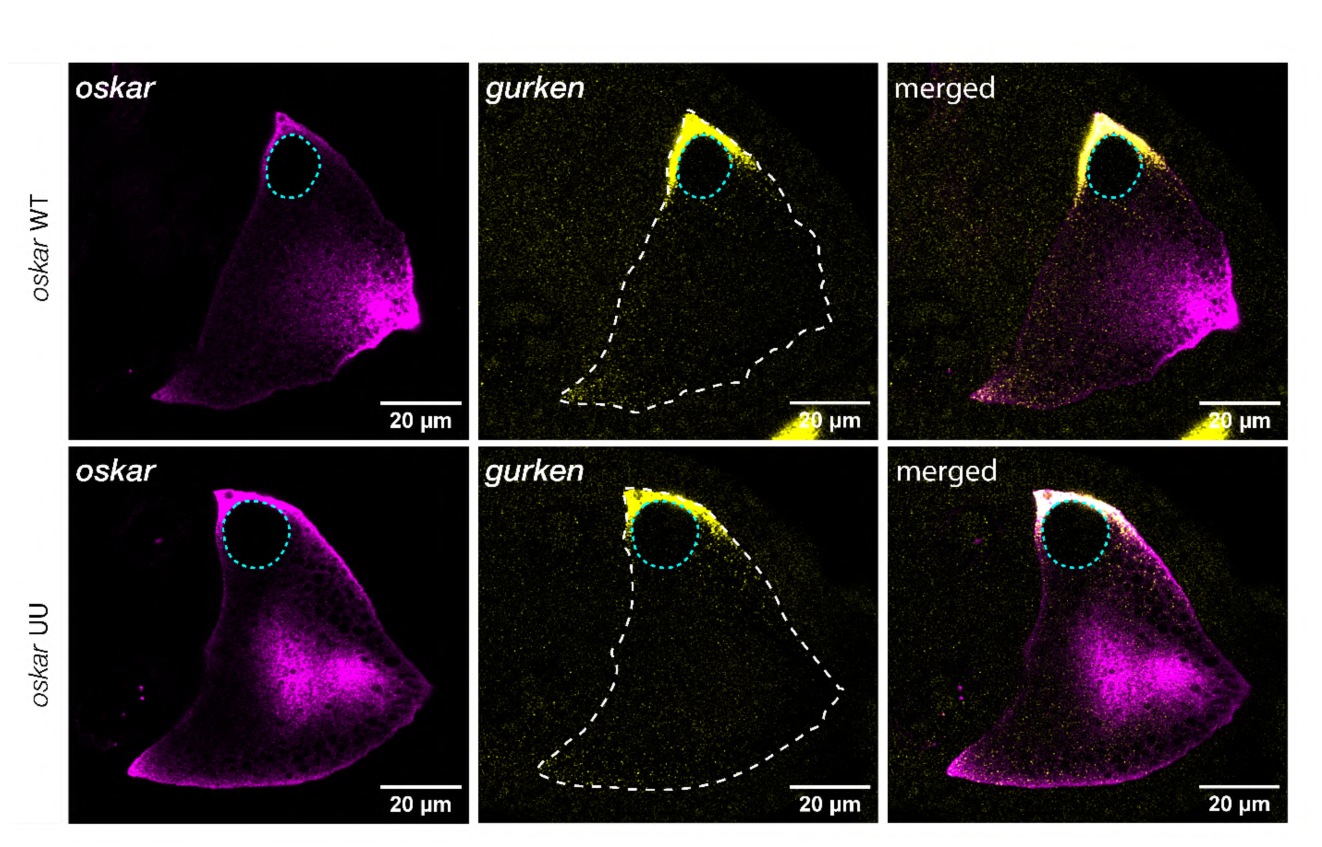
*oskar* (magenta) and *gurken* (yellow) smFISH images in *oskar* WT and UU egg chambers indicating that oocyte polarity is intact in both genotypes. Also note the anterior-lateral position of the nucleus (cyan dotted line) which is another marker of oocyte polarity.

**Extended Data Fig. 2.**
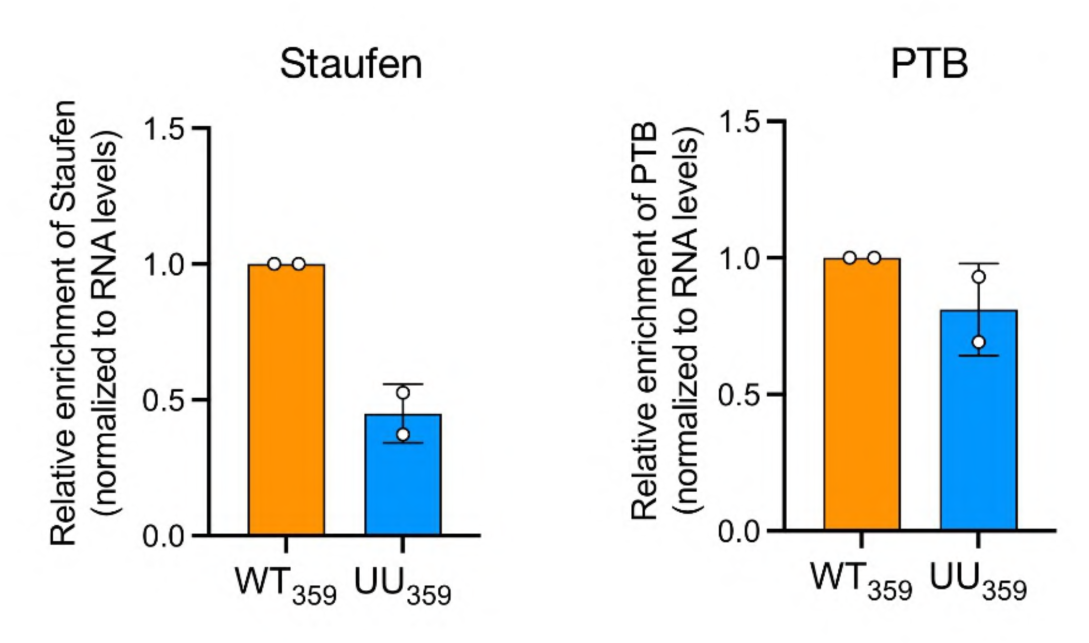
Quantification of the Staufen and PTB levels associated with WT_359_ and UU_359_ RNAs carried out using western blots from two independent replicates after normalization with the recovered RNA level. Error bars represent SD.

**Extended Data Fig. 3.**
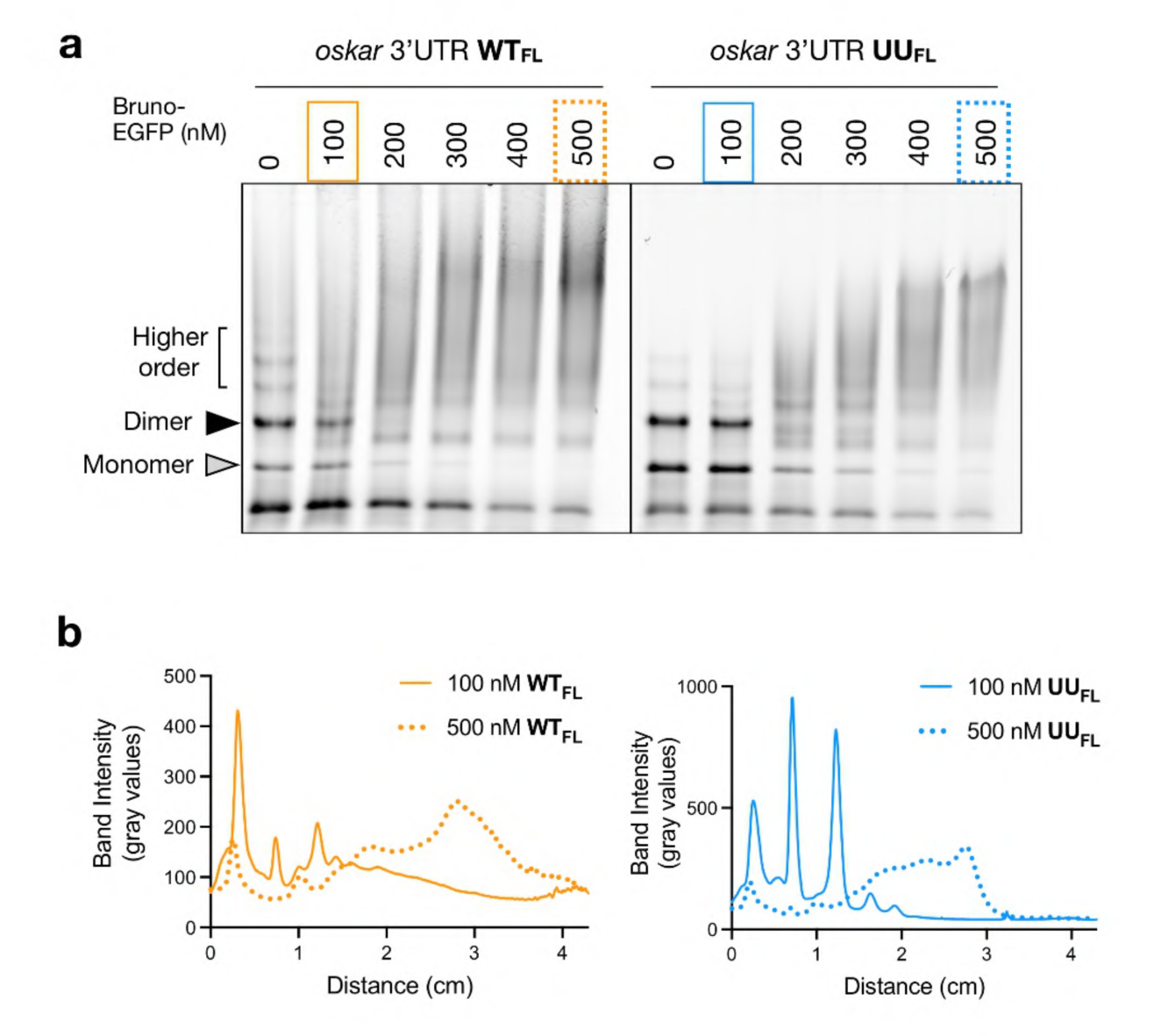
**a,** EMSA of the atto633-labeled full length (FL) *oskar* 3’UTR with indicated concentrations of Bruno-EGFP. The WT and UU samples are from different regions of the same gel, as indicated by the black line. **b,** Lane intensity profiles (from bottom to top of the gel) of the respective RNAs at indicated concentrations of Bruno-EGFP.

**Extended Data Fig. 4.**
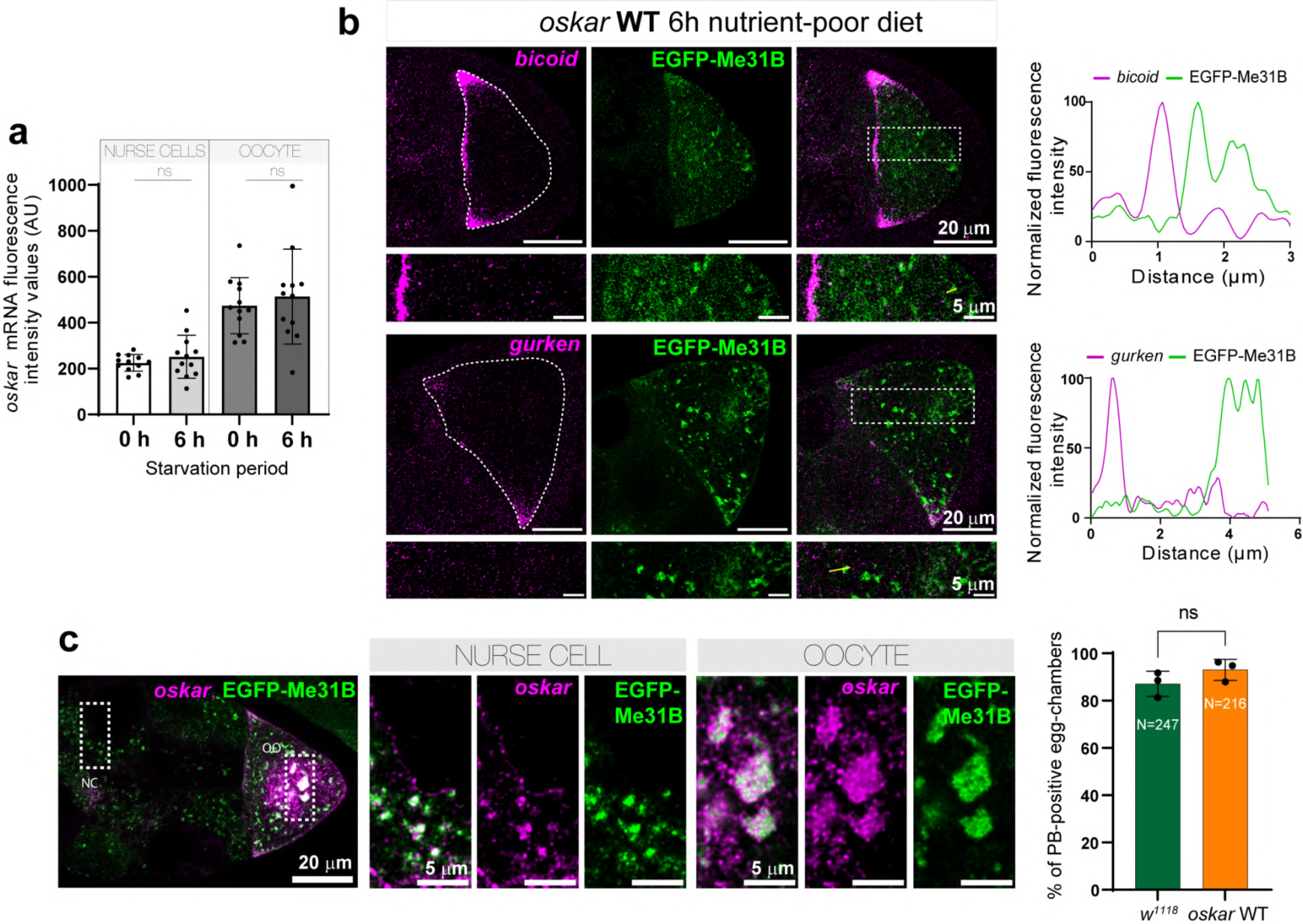
**a,** Quantified intensities of *oskar* smFISH signal under nutrient-rich (0 h) and nutrient-poor (6 h) conditions in nurse cells and oocytes. **b,** smFISH detection of *bicoid* mRNA (magenta; top panel) or *gurken* mRNA (magenta; bottom panel) in EGFP:Me31B (green) expressing *oskar* WT flies (in absence of endogenous *oskar* RNA). Boxed areas are enlarged below the images. Representative line profiles (yellow) are plotted on the right. **c,** Co-detection of *oskar* WT (magenta) and EGFP-Me31B (green) under nutrient-poor conditions. Boxed areas are enlarged on the right. The graph on the right shows the percentage of egg chambers forming P-bodies in wild-type (*w*^1118^) and transgenic *oskar WT* flies (in absence of endogenous *oskar* RNA). Three biological replicates were analyzed and egg chambers at stages 6-9 were scored. Unpaired two-tailed t-test was used for the statistical analysis and mean ±SD was plotted (values of significance: *,p ≤ 0.05; and if not as ns; N = number of egg chambers analyzed).

## Methods

### Fly stocks & fly husbandry

The following *Drosophila melanogaster* wild-type fly stocks were used: *w*^1118^ and *Oregon-R* (for the RNA affinity pull-down assay). To generate the *oskar* RNA null background the following fly stocks were used: *oskar^A^*^87 20^, *Df(3R)p^XT^*^103 15^ and *oskar^attP,3P^*^3*-GFP* 46^. EGFP:Me31B fly line^47^ was used to visualize P-bodies in the *Drosophila* germline. *oskar* WT, *oskar* UU transgenes were expressed in the *Drosophila* germline under control of the pUASP promoter using the *oskar*-GAL4 driver. The Bruno KI-EGFP line was a gift of Akira Nakamura^27^.

All fly lines were maintained at 18°C or 25°C in vials or bottles on standard food (corn-based medium). 3-6 day old female flies with typically half as many male flies were transferred to vials with fresh yeast 24 h before ovary dissection. For nutrient deprivation, flies were kept on standard food supplemented with dry yeast granules for 24 h at 25°C (nutrient-rich conditions) and transferred to fresh standard food without dry yeast (nutrient-poor conditions) and kept for 6 hours at 25°C. Ovaries were then dissected and processed for smFISH and/or antibody staining.

### Single molecule fluorescent *in situ* hybridization (smFISH) and immunostaining

Oligonucleotide probes (18-22 nt) spanning the coding sequence and the 3’UTR of *oskar* mRNA were labeled using atto633 (Atto-Tec GmbH) as described previously^48^. Freshly dissected *Drosophila* ovaries were fixed with 2% paraformaldehyde (PFA) in PBS with 0.05% Triton X-100 (Sigma) followed by two rounds of 10 min washes in PBS supplemented with 0.1% Triton X-100 (PBT). Ovaries were then pre-hybridized in hybridization buffer (HyB: 2x SSC, 1 mM EDTA, 1 v/v % Triton X-100, 15 v/v % ethylene carbonate, 50 μg/ml heparin, 100 μg/ml single stranded DNA from salmon testes) for 10 min at 42°C. Hybridization was performed at the same temperature in HyB containing probe mix (2–3 nM per probe) for 2-3 h. After hybridization, the following 10 min × 5-step washing protocol (a–d) was performed (a-HyB at 42°C, b-1:1 HyB + PBT at 42°C, c-PBT at 42°C, d-pre-warmed PBT (42°C) at room temperature, e-PBT at room temperature). After washes, embedding or immunostaining was performed. To detect EGFP-tagged proteins, native fluorescence of EGFP was visualized. To detect endogenous Me31B, immunostaining was applied following smFISH. Ovaries were incubated overnight with mouse α-Me31B antibody (1:200; from Nakamura) in PBT supplemented with 1xβ-casein at +4°C, followed by three 10 min washes in PBT at room temperature. The ovaries were then incubated with secondary antibody (1:750; anti-mouse AlexaFluor633 IgG (H+L) Highly Cross-Adsorbed) in 1xβ-casein/PBT for 2 h at room temperature followed by three 10 min washes in PBT at room temperature. Ovaries were mounted in 25–50 μl 80% TDE or in 85% glycerol supplemented with 2% propyl gallate. Counterstaining with DAPI was done to visualize the nuclei.

### Microscopy

All images were acquired using a Leica SP8 TCS X confocal laser scanning microscope with a HC PL APO 63x/1.30 Glycerol CORR CS2 glycerol-immersion objective. Images were deconvolved with the Huygens Essentials software or on the fly using the Leica Lightning module. For *in vitro* phase separation experiments, images were acquired with an HC PL APO 40x/1.10 W CORR CS2 water-immersion objective without any further deconvolution.

### *In vitro* transcription, fluorescent labeling and visualization

DNA templates for the IVT reactions were prepared by PCR using T7-forward primer and gene specific reverse primers and extracted by gel purification. IVT was performed with MegaShortScript T7 Kit (Invitrogen) for the 359 and 67 nucleotide long fragments and subsequent RNA purification was performed according to manufacturer protocol. For the full length *oskar* 3’UTR (WT and UU), MegaScript T7 Kit (Invitrogen) was used and RNA recovered by lithium chloride precipitation.

For fluorescent labeling, the IVT reaction was doped with 5-amino-allyl UTP (Biotium) at 1:10 (aminoallyl-UTP: UTP) followed by incubation of purified transcripts with 3-fold molar excess of atto633 NHS-ester (Atto-Tec GmbH) in 0.1 M NaHCO_3_ at RT for 2 h, protected from light. The RNA was extracted using absolute ethanol and sodium acetate, pH 5.5 precipitation at −20°C for at least 1 hour and dissolved in ultrapure water (Invitrogen). Transcript quality was checked with SYBR Safe stain (473 nm) and fluorescent imaging (635 nm) in a Typhoon biomolecular imager.

For visualization of RNA under indicated conditions, the labeled RNA was incubated at RT for 15 min in the indicated buffers and electrophoresed on a 0.8-1% agarose gel (0.5x TBE) run at 100V at 4°C. For the 67 nt. OES fragments, native 6% acrylamide gels (Novex™ TBE-Urea Gels, Invitrogen) were used. The following buffer conditions were used : stringent zero salt buffer (20 mM Tris-HCl pH 7.5, 5% glycerol, 0.5 mM TCEP); EMSA buffer (20 mM Tris-HCl pH 7.5, 150 mM NaCl, 2 mM MgCl_2_, 5% glycerol, 0.5 mM TCEP); high salt buffer (20 mM Tris-HCl pH 7.5, 300 mM NaCl, 5 mM MgCl_2_, 5% glycerol, 0.5 mM TCEP).

### 3’ end labeling of RNA with Biotin

The *in vitro* transcribed RNAs were 3’ biotinylated using pCp-Biotin (Jena Biosciences). Briefly, 2 µM transcript was incubated with 10-fold molar excess of pCp-Biotin in T4 RNA Ligase buffer, 10% DMSO, 1mM ATP, 16% PEG-8000 and 1 unit/µl T4 RNA Ligase enzyme at 16°C overnight. The RNA was recovered by acidic phenol-chloroform extraction and reconstituted with ultrapure water.

### RNA affinity capture assay

Equimolar amounts of 3’-biotinylated RNA were added to ovary Lysis Buffer (5 mM Hepes-NaOH pH 7.5, 1 mM MgCl2, 50 mM KCl, 25 mM sucrose, 0.5% NP-40, 50-75U of ribolock (Thermo Fisher Scientific), 0.5% Triton-X 100 (Sigma), 1 tablet of EDTA-free Protease Inhibitor Cocktail (Merck), 10 mM DTT) and incubated with 25 µl of pre-washed paramagnetic streptavidin beads (Dynabeads MyOne Streptavidin C1 beads, Invitrogen) at 4°C for 2 hours.

Meanwhile, ovary lysate was prepared. To obtain an optimal amount of ovary lysate for the pull-down assay, 30–40 g of *Oregon-R* flies was harvested^24^. 3–4 ml of harvested ovaries were lysed in the Lysis buffer using a Dounce homogenizer. The lysate was clarified and incubated with Avidin-agarose beads at 1/50^th^ of the lysate volume for 30 min at 4°C to remove endogenous biotinylated proteins. After removing the Avidin-agarose by mild centrifugation, the lysate volume was adjusted to 12 ml using the lysis buffer and the lysate divided into 6 equal halves for the no RNA control, WT_359_ and UU_359_ conditions in duplicates. The respective bead-conjugated RNA was added to the lysate and incubated at 4°C for 1 h in a nutator. Subsequently, 3 × 10 min washes were performed in the wash buffer (25 mM Hepes-NaOH pH 7.5, 1 mM MgCl2, 150 mM KCl, 25 mM sucrose, 0.5% NP-40). In the final wash, 80% of the sample was eluted in Laemmli buffer supplemented with β-mercaptoethanol for western blotting and remaining 20% was used for RNA extraction using Trizol LS for qPCR-based detection of pulled-down RNA levels.

### Western blotting

Western blotting was performed using the following primary antibodies: rabbit α-Bruno (1:1000, in-house), rabbit α-Staufen (1:5000, in-house), rabbit α-PTB (1:2000, in-house), rabbit α-Egalitarian (1:2500)^49^. Anti-rabbit secondary antibody conjugated with horseradish peroxidase (GE Healthcare) was used for chemiluminescence-based detection.

### RNA extraction, cDNA synthesis and qPCR

SuperScript III First-Strand Synthesis System SuperMix (Invitrogen) was used for first-strand cDNA synthesis from isolated RNA using the manufacturer’s instructions. Random hexamers were used for cDNA synthesis and the following gene specific primers for detecting *oskar* 359 fragment in the affinity-purified samples : 5’-GCGCGATTTTCGTCTTTCTGTTTC-3’ (forward) and 5’-GTAGCACAGTGTAGAATTCTGGCG-3’ (reverse).

### Protein purification

The pCoofy63-BrunoFL-EGFP (6xHis-SumoStar-BrunoFL-EGFP-TwinStrep) construct was used for expression and purification of Bruno as described previously^8^. Briefly, half a liter of Sf-21 insect cells were infected at 0.5-0.7×10^6^ cell/ml density with the recombinant baculovirus stock at a ratio of 1:100 and cells harvested 72 hours post-infection. The cell pellet was flash-frozen and stored at −80°C. For purification, the pellet was resuspended in lysis buffer (20 mM Tris-HCl pH 7.5, 500 mM NaCl, 1 mM EDTA supplemented with 0.01% TritonX-100, 1x tablet of Complete Mini Protease Inhibitor cocktail (Roche), 2 mM MgCl_2_) for 10 min on ice, followed by digestion with Benzonase (Sigma) for digestion of RNA/DNA and lysed using a microfluidizer. The lysate was clarified by centrifugation at 16,000 x g at 4°C for 20 min and the protein affinity-purified using a StrepTrap HP column by the C-terminal TwinStrep tag. The column was washed with 5-6 column volumes of wash buffer (20 mM Tris-HCl pH 7.5, 500 mM NaCl, 1 mM EDTA) and eluted in wash buffer supplemented with 2.5 mM desthiobiotin (Sigma). Protein-enriched fractions were pooled and dialysed overnight to remove EDTA and desthiobiotin. The following day the protein was concentrated to 5 ml and subjected to size exclusion chromatography using a HiLoad 16/600 Superdex 200 pg column in storage buffer (20 mM Tris-HCl pH 7.5, 300 mM NaCl, 2 mM MgCl_2_, 5% glycerol, 0.5 mM TCEP). Desired fractions were collected and concentrated using 50 kDa MWCO concentrators (Amicon), aliquoted and flash-frozen for storage at - 80°C. Importantly, during concentrating the protein post-size exclusion chromatography, the sample was frequently checked under a fluorescence microscope to make sure that no phase separation occurs.

### Electrophoretic Mobility Shift assays

EMSA was carried out as described previously^8, 26^. 50 nM of atto633-labeled *oskar* RNA constructs was incubated with increasing concentrations of BrunoFL-EGFP for 20 min at RT in EMSA buffer (20 mM Tris-HCl pH 7.5, 150 mM NaCl, 2 mM MgCl_2_, 5% glycerol, 0.5 mM TCEP) and resolved on a cold 0.8% agarose gel (0.5X TBE) run at 100V at 4°C. Imaging of the gel was performed in a Typhoon biomolecular imager.

### *In vitro* phase separation assays

All *in vitro* phase separation assays were carried out in assay buffer (20 mM Tris-HCl pH 7.5, 150 mM NaCl, 2 mM MgCl_2_, 5% glycerol, 0.5 mM TCEP) and reactions spotted on 96-well non-binding µclear plates (Greiner Bio-One) for microscopy. A frozen aliquot of the protein was thawed and centrifuged at high speed to get rid of pre-formed aggregates, protein concentration measured and reaction assembled with or without 100 nM of atto633-labeled RNAs. The 6xHis-SumoStar tag was maintained during the experiments. Details of individual experiments are indicated in the respective figure legends.

### Image analysis

Analysis and processing of acquired images were performed using Fiji. Maximum intensity projections of image stacks, histogram generation as well as lane profiles of the agarose gels and EMSAs were also calculated with Fiji.

#### Quantification of fluorescence intensity

To quantify *oskar* mRNA, 2D sum projections were prepared for the oocyte and nurse cell compartments using on average 20 z-planes with a z-step size of 0.9 μm. For a region of interest, raw integrated density of smFISH signal was measured and intensities were calculated as:

I = raw integrated density/(area(μm^2^) x z-stack size (μm)). All representative images are 2D maximum projections of on average 2 to 3 planes of acquired xyz-stack unless indicated otherwise.

#### Quantification of oskar localization

*oskar* mRNA distribution was analyzed using the CortAnalyis Fiji plugin as previously described elsewhere^8,46,50^.

#### Quantification of oskar partition coefficient in the oocyte

Intensity-based segmentation of the granules was carried out based on the *oskar* smFISH signal for the *oskar* WT and *oskar* UU genotypes. After trials with several segmentation algorithms, the Triangles algorithm could reliably distinguish granules from the diffuse RNA signal in the dilute phase in case of *oskar* UU. Partition coefficient was calculated as the ratio of the mean intensity inside granules to that of the oocyte cytoplasm^8^.

#### Quantification of condensate parameters

For quantification of particle size and area occupied, the RNA channel was filtered with a Gaussian filter of radius of 1 pixel, particles segmented using the Triangles algorithm and areas calculated using the particle analysis in Fiji.

### Statistical analysis

For all quantifications, statistical analyses were performed and the data plotted using Prism 9. P-values were calculated using unpaired two-tailed t-tests. In the figure, * = p< 0.05, ** = p< 0.01, *** = p< 0.001, **** = p< 0.0001.

